# Task-Evoked Functional Connectivity Does Not Explain Functional Connectivity Differences Between Rest and Task Conditions

**DOI:** 10.1101/252759

**Authors:** Lauren K. Lynch, Kun-Han Lu, Haiguang Wen, Yizhen Zhang, Andrew J. Saykin, Zhongming Liu

**Author notes:** Correspondence Zhongming Liu, PhD Assistant Professor of Biomedical Engineering Assistant Professor of Electrical and Computer Engineering College of Engineering, Purdue University 206 S. Martin Jischke Dr., West Lafayette, IN 47907, USA.

## Abstract

During complex tasks, patterns of functional connectivity (FC) differ from those in the resting state. What accounts for such differences remains unclear. Brain activity during a task reflects an unknown mixture of spontaneous activity and task-evoked responses. The difference in FC between a task state and resting state may reflect not only task-evoked connectivity, but also changes in spontaneously emerging networks. Here, we characterized the difference in apparent functional connectivity between the resting state and when human subjects were watching a naturalistic movie. Such differences were marginally (3-15%) explained by the task-evoked networks directly involved in processing the movie content, but mostly attributable to changes in spontaneous networks driven by ongoing activity during the task. The execution of the task reduced the correlations in ongoing activity among different cortical networks, especially between the visual and non-visual sensory cortices. Our results suggest that the interaction between spontaneous and task-evoked activities is not mutually independent or linearly additive, and that engaging in a task may suppress ongoing activity.

## Introduction

Functional connectivity (FC) captures the correlation of different networks or regions of the brain. Its structure and dynamics have been useful in characterizing the brain’s functional organization. Patterns of FC are similar across distinct states of consciousness (Horovitz, et al., 2008; Vincent, et al., 2007), and they are also largely conserved during the performance of various tasks (Arfanakis, et al., 2000; Cole, et al., 2014; Fair, et al., 2007; Gratton, et al., 2016; Harrison, et al., 2008; Krienen, et al., 2014). However, increasing evidence suggests that FC is altered within and between brain states (Buckner, et al., 2013; Chang and Glover, 2010; Hutchison, et al., 2013; Mennes, et al., 2013; Rehme, et al., 2013; Sepulcre, et al., 2010; Van Dijk, et al., 2010; Wong, et al., 2013). It leads to the potential use of FC as a network signature of how the brain engages itself in various behavioral or cognitive tasks, e.g. watching a movie. In fact, FC signatures have been used to accurately classify a multitude of brain states (Gonzalez-Castillo, et al., 2015), leveraging this notion.

During a task, brain activity measurements reflect a mixture of spontaneous and evoked activities. Disentangling their differential contributions to the pattern of apparent FC is essential to proper interpretation of any FC difference between a task and resting-state, or between different tasks. If the task-dependent FC is due to the task-evoked activity, its pattern reflects the network interactions directly involved in information processing for task execution. If the task-dependent FC is attributed to ongoing activity, its pattern is driven by the brain’s functional re-organization or adaption to facilitate the task. Alternatively, evoked activity may interact with spontaneous activity. As such, the task-dependent FC should reflect correlational changes in both task-evoked networks and spontaneously emerging networks.

There is a lack of consensus on the relationship between evoked and ongoing activities. Some prior studies suggest that task-evoked activity is independent from spontaneous neural processes (Arieli, et al., 1996; Mäkinen, et al., 2005; Tsodyks, et al., 1999). Initial evidence has led to the notion that spontaneous and evoked processes linearly sum to yield the activity observed during a task (Arieli, et al., 1996; Azouz and Gray, 1999; Becker, et al., 2011; Fox, et al., 2006; Saka, et al., 2010). There are, however, other reports to the contrary. Using electrophysiology, several groups have shown a reduction in neural variability following the onset of a stimulus, suggesting that the task suppresses ongoing activity during the task (Borg-Graham, et al., 1998; Churchland, et al., 2010; Finn, et al., 2007; Oram, 2011; Ponce-Alvarez, et al., 2013). Using fMRI, He et al. (2013) also found a negative interaction between spontaneous activity and task-evoked activity during a visual attention task. However, how (and whether) such an interaction may occur with respect to functional connectivity has not been fully investigated.

Prior studies have established some valuable analysis methods to address this question. Simony et al. proposed the use of inter-subject functional connectivity (ISFC) during sustained and natural stimulation to extract task-evoked networks without contributions from ongoing activity or non-neuronal noise (Simony, et al., 2016). For any given pair of regions, cross-correlation between one subject’s time series in one region with the mean time series from all other subjects in the other region was only attributable to task-evoked activity. This technique builds off of the Hasson, et al. (2004) study, which showed that natural stimulation gave rise to reliable responses reproducible across individuals. Like ISFC, a similar strategy is to assess the inter-regional correlation across different sessions of the same stimuli for the same subject (Lu, et al., 2016; Wilf, et al., 2017), while further discounting the variation across subjects.

Using this strategy in this study, we sought to examine whether task-evoked networks were additive to spontaneous networks, and if they were able to explain the change in FC during movie watching relative to the resting state (or the “task-rest FC” difference for simplicity). To address these questions, we began with examining the seed-based correlations for exploratory analysis, and subsequently performed systematic analysis of functional connectivity among brain parcels or networks.

## Materials and Methods

### Subjects

Thirteen healthy volunteers (20 – 31 years old, 8 females, 12 right-handed, normal or corrected to normal vision) participated in this study in accordance with a protocol approved by the Institutional Review Board at Purdue University. Three subjects were excluded because they either were self-reported to fall asleep or had excessive head motion during the experiment.

### Experimental design

Each of the remaining 10 subjects (20-31 years old, 6 females, 9 right-handed) underwent four fMRI sessions with two conditions. Two sessions were obtained in the eyes-closed resting state, and the other two sessions occurred during free-viewing of an identical movie clip (The Good, the Bad, and the Ugly, 1966, from 162:54 to 168:33 min. in the film), as used in prior studies (Hasson, et al., 2004; Lu, et al., 2016). The visual stimulus was presented using the MATLAB Psychophysics Toolbox (Brainard, 1997; Pelli, 1997); it was delivered to the subjects through a binocular goggle system (NordicNeuroLab, Norway) mounted on the head coil. The display resolution was 800×600; through the goggle system, the visual field covered by the movie was about 26.9°×20.3°. No sound was presented during the movie. Each movie-stimulation session began with a blank gray screen presented for 42 s, followed by the movie presented for 5 min and 37 s, and ended with the blank screen again for 30 s. The resting-state sessions had the same duration as the movie-stimulation sessions. The session order was randomized and counterbalanced across subjects. For simplicity, hereafter the resting-state and movie-stimulation sessions were referred to as the “rest” and “task” conditions, following the general notions in a broader context (Cole, et al., 2014).

### Data acquisition

Whole-brain structural and functional MRI images were acquired using a 3-Tesla Signa HDx MRI system (General Electric Health Care, Milwaukee, USA). As described previously (Marussich, et al., 2017), the fMRI data were acquired using a single-shot, gradient-recalled (GRE) echo- planar imaging (EPI) sequence (38 interleaved axial slices with 3.5mm thickness and 3.5 × 3.5 mm^2^ in-plane resolution, TR=2000 ms, TE=35 ms, flip angle=78°, field of view=22×22 cm^2^). T1-weighted anatomical images covering the whole head were acquired with a spoiled gradient recalled acquisition (SPGR) sequence (1×1×1mm^3^ voxel size, TR/TE=5.7/2ms, flip angle=12°). A 16-channel receive-only phase array coil (NOVA Medical, Wilmington, USA) was used for image acquisition.

### Pre-processing

Pre-processing of the fMRI images was carried out with a combination of AFNI (v17.0.01) (Cox, 1996), FSL (v5.0.8) (Smith, et al., 2004), and MATLAB 2017A (Mathworks, Natick, MA). T_1_-weighted anatomical images were non-linearly registered to the Montreal Neurological Institute (MNI) brain template using a combination of *flirt* and *fnirt* in FSL (Smith, et al., 2004). T_2_*-weighted functional image time series were corrected for slice time variations using *slicetimer* in FSL, co-registered to the first volume within each series to account for head motion using *mcflirt* in FSL, restricted to within-brain tissues using *3dcalc* in AFNI (Cox, 1996), aligned to the T_1_-weighted structural MRI using FSL’s Boundary Based-Registration (BBR) function (Greve and Fischl, 2009), and registered to the MNI space with 3-mm isotropic voxels using *applywarp* in FSL. The first six volumes in the fMRI data were discarded to avoid any pre-steady-state longitudinal magnetization.

The remaining pre-processing steps were conducted in MATLAB using in-house code. For the task sessions, we only analyzed the fMRI data during the movie while excluding any transient fMRI response during the first few seconds since the start of the movie. Thus, we excluded the first eight seconds and the last fourteen seconds of the movie. For each session and each voxel, the voxel time series was detrended by regressing out a third-order polynomial function that modeled the slow trend; the detrended signal was bandpass filtered (0.0001 - 0.1 Hz). Spatial smoothing was applied by using a Gaussian kernel (FWHM=6 mm), and the spatially smoothed voxel time series were demeaned and normalized to unit variance.

### Seed-based functional connectivity in rest versus task

We first explored the difference in seed-based correlation patterns between the resting state and the task state. For this purpose, seed voxels were selected from the primary visual cortex (V1), higher visual areas (HV), precuneus (PCu), and primary motor cortex (M1); each of these regions of interest was defined using an atlas from an independent study (Shirer, et al., 2012)^1^. The MNI coordinates of these seed regions were (0, −54, 30) for PCu, (0, −87, 9) for V1, (48, −78, 0) for HV, and (3, −18, 57) for M1. These seed locations were chosen because they are representative of major functional systems activated by visual (V1 and higher visual areas) or motor tasks (M1), or deactivated by cognitive tasks (PCu as a part of the default-mode network).

Within either a rest or task session, the correlation between the seed voxel’s time series and every other voxel’s time series was calculated (after global signal regression), and the correlation coefficient was converted to a z-score using the Fisher’s transform. The voxel-wise z-score was averaged across all rest (or task) sessions from all subjects. The significance of the mean z-score (against zero) was evaluated by using one-sample t-test (df = 19) corrected for multiple comparisons at the false discovery rate (FDR) q<0.03. The above analysis was performed separately for the rest and task conditions.

To determine the task-rest FC difference, the mean z-score of the movie sessions was then compared to the mean z-score of the resting-state sessions using a paired t-test (df = 19, p<0.03, uncorrected). Then, to determine the task-evoked FC, the seed voxel’s time series in session 1 was cross-correlated with the time series of all voxels in session 2 for each subject; the resulting Pearson correlation values were then z-transformed. This process was repeated was repeated using seed voxels in session 2 with cross-correlations to all voxels in session 1. To determine the statistical significance of the results, the mean z-score was compared to zero using one-sample t-tests for the task-evoked connectivity (df = 19, q<0.05, FDR corrected).

### Whole Brain Functional Connectivity

To compare the task-rest FC difference to the task-evoked FC in a systematic manner encompassing the whole brain, neural activity was decomposed into smaller networks and/or regions using three different methods: 1) using a 17-network atlas (Yeo, et al., 2011), 2) via networks obtained using spatial independent component analysis (ICA), and 3) using a fine-grained, 246-region functional atlas (the Brainnetome Atlas, v1.0) (Fan, et al., 2016). The 17-network atlas was from http://surfer.nmr.mgh.harvard.edu/fswiki/CorticalParcellation_Yeo2011^2^ and the 246-region Brainnetome Atlas was obtained from http://atlas.brainnetome.org/^3^. Using the 17-network and 246-functional parcellation atlases, the mean intensity of brain regions over time was regressed from the signal.

Group spatial ICA using the Infomax algorithm (Bell and Sejnowski, 1995) was applied to data after two additional processing steps. Prior to ICA, the data was concatenated across all subjects, sessions, and conditions; principal component analysis (PCA) was applied to the data such that 95% of the variance was retained. After ICA was applied to this result, 30 independent components were obtained; of those, 24 networks corresponded to canonical resting-state networks (RSNs) (Beckmann, et al., 2005; De Luca, et al., 2006; Power, et al., 2011). One component with a global pattern was excluded from the analysis. Then each session’s time course was obtained by regressing the group spatial map into the session’s 4D dataset.

Large-scale FC was assessed within the resting-state and within the movie task (“mixed” FC). To create the within-session resting-state and movie FC, the correlations between each pair of networks or regions calculated based on based on their corresponding time series, and then the correlation coefficient was converted to the z-score. Significant correlations were identified using one-sample t-tests for each pair of regions or networks in each condition (df = 19), FDR-corrected at q < 0.03. Then, task-rest FC differences were evaluated by subtracting the resting-state z-scores from the movie z-scores for each network pair. Significant differences were evaluated using paired t-tests (df = 19), FDR corrected at q< 0.03. Finally, to obtain the task-evoked FC, the cross-correlations between each network/region’s mean time series in session 1 and the mean time series in session 2 were determined and z-transformed. To create symmetric mean FC matrices that would also be comparable to the other profiles, we included the transposes of the task-evoked FC matrices. We then evaluated significant correlations using one-sample t-tests (df = 19), FDR-corrected at q < 0.03. Because our focus was on the functional connectivity between regions or networks, we ignored the correlation within the exact same region or network itself in our analysis.

In order to further characterize the similarity of the FC profiles, we performed a session-wise cross-correlation analysis of the FC matrices. Spatial cross-correlations between the resting-state FC matrices and the movie FC matrices were calculated for each pair of sessions (e.g. Subject 1 Rest Session 1 with Subject 1 Movie Session 1, and so on). Lower triangular matrices (from one element below the diagonal) were used in these correlation calculations to represent only unique, meaningful information from the symmetric matrices. Then, spatial cross-correlations were calculated between the task-rest FC difference matrices and the task-evoked FC matrices.

In this case, the lower triangular content of the first session’s FC difference (e.g. Subject 1 Movie Session 1 – Subject 1 Rest Session 1, a symmetric matrix) was correlated with that subject’s lower triangular task-evoked FC (e.g. Subject 1 Movie Session 1 with Subject 1 Movie Session 2, not a symmetric matrix), and the second session’s difference (e.g. Subject 1 Movie Session 2 – Subject 1 Rest Session 2) was correlated with that subject’s upper triangular task-evoked FC (e.g. Subject 1 Movie Session 1 with Subject 1 Movie Session 2). The reasons for this were two-fold: first, we performed this at the subject level to maximize the amount of information obtained from individual subjects; secondly, because the cross-correlation of one session’s information with that of a second session does not yield a symmetric matrix, taking into account the upper triangular/lower triangular transpose of this information (e.g. the cross-correlation of second session’s information with the first session’s information) reveals the full spectrum of connectivity information.

### Comparing significant task-rest FC differences with task-evoked FC

The specific functional connectivity implicated in the task-rest FC difference and the task-evoked FC were investigated using the fine-grained, 246-region parcellation’s information. To test the significance of the functional connectivity between each pair of regions and/or networks, the average z-score was compared against zero by performing one- sample t-test on the z-score of every pair regions (q < 0.03, FDR corrected).

### Explaining the task-rest FC difference with task-evoked FC

To determine the extent to which the task-evoked FC explains the task-rest FC difference, the task-evoked FC matrices were linearly regressed into the task-rest FC difference matrices at the session-level. This was performed separately using 1) the Yeo et al. 17-network atlas (2011), 2) the previously obtained 24 spatial ICs, and 3) the 246-region Brainnetome Atlas (Fan, et al., 2016). As was conducted in the Whole Brain Functional Connectivity section, the lower triangular content of the first session’s FC difference was regressed into that subject’s lower triangular task-evoked FC, and the second session’s lower triangular FC difference was regressed into that subject’s upper triangular task-evoked FC. After obtaining a regression coefficient for each session, the estimated task-rest FC difference was obtained by multiplying the calculated coefficients with the task-evoked FC. Then, the variance of this estimated task-rest FC difference was divided by the variance of the measured task-rest FC difference to yield the percentage of the task-rest FC difference that was explained by the task-evoked FC.

## Results

### Seed-based FC Distributions

Seed voxels from the PCu, V1, HV, and M1 were used to assess voxel-wise FC at rest, voxel-wise FC during the movie task (i.e. the “mixed” FC), the difference between these two states, and the task-evoked FC (Fig. 1, findings projected onto the surface).

**Figure 1.**
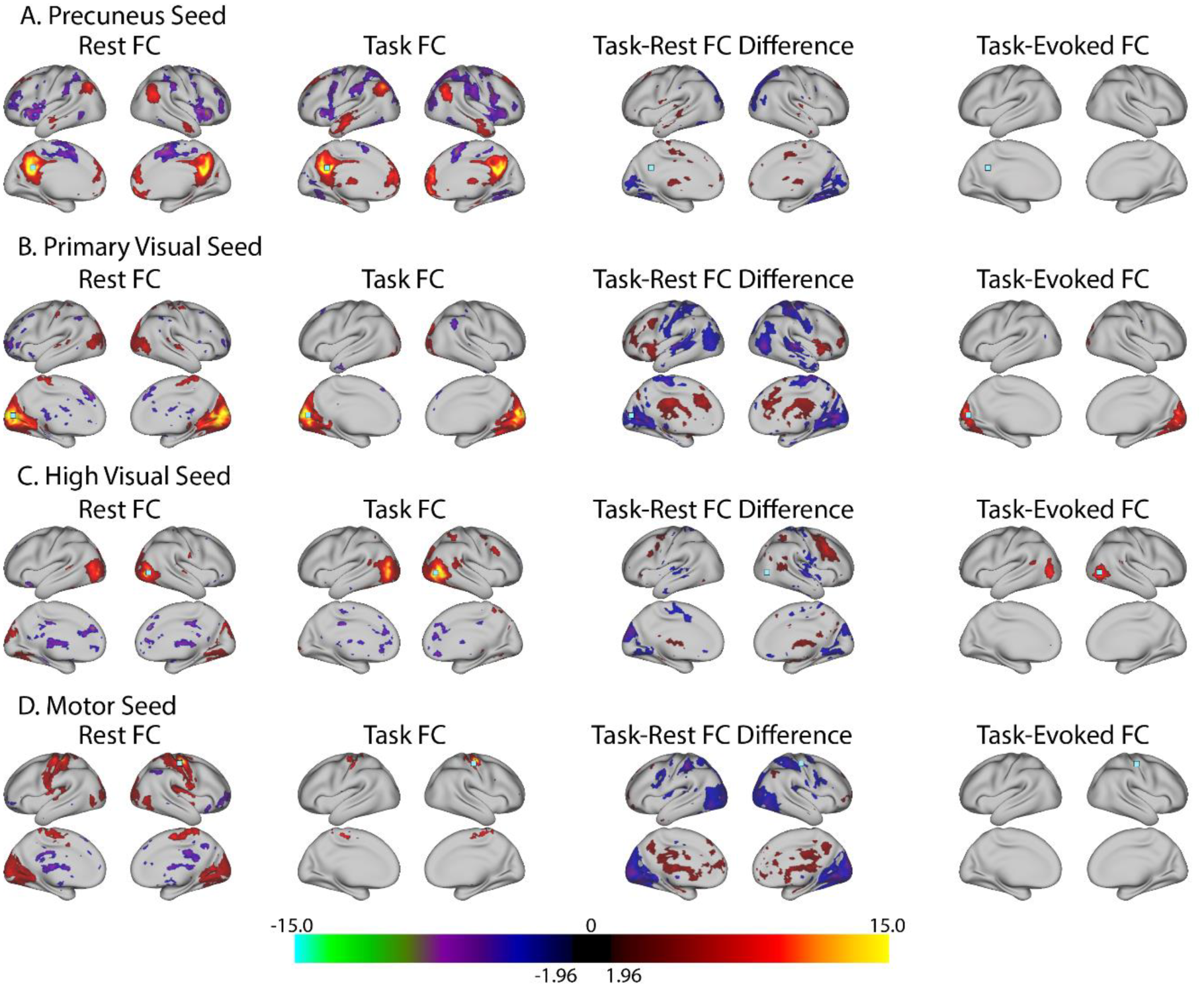
Seed-based FC Findings. Seed-based functional connectivity (p<0.03, FDR-corrected for all except for within-session differences) using a seed in A) PCu, B) V1C) HV, and D) M1. Each panel shows the result for within-session FC during eyes-closed resting-state (left), within-session FC during the movie task (left middle), the within session FC difference during the movie relative to rest (right middle), and the task-evoked FC computed using inter-session correlations (right). The seed voxel is shown as a light blue square in each image. The color bar indicates t-values.

FC patterns in the resting-state and during the movie were mostly consistent among the four seeds, but there were some differences between the two conditions. Although the PCu seed exhibited similar, positive distributions in both conditions, the anti-correlated voxels were more widespread during the movie task (Fig. 1A, far left and left middle columns). In the resting-state, the V1 seed (Fig. 1B, far left and left middle columns) was coupled not only to higher visual areas, but also to the superior/medial motor cortex; during the task, the broad primary visual cortex indeed was produced, but no coupling to other networks was observed. In addition, the HV seed was more connected to more medial visual areas (e.g. fusiform gyri) during resting-state (Fig. 1C, far left and left middle columns). Finally, the distribution of the FC from the M1 seed elicited visual networks at rest but was more narrowly confined during the movie task (Fig. 1D, far left and left middle columns).

The task-evoked and task-rest FC difference distributions were largely very different. Task-evoked FC was observed using the V1 and HV seeds but not using the PCu and M1 seeds, in line with the findings of others (Kim, et al., 2017; Wilf, et al., 2017). The positively connected voxels to V1 in the task-evoked FC (Fig. 1B, far right column), were, in fact, more weakly connected to the seed during the movie than at rest (Fig. 1B, right middle column). Moreover, using the HV seed, the positively connected voxels arising from task-evoked activity (Fig. 1C, far right column) were not significantly different between the movie and the task (Fig. 1C, right middle column) despite qualitatively appearing stronger during the movie (Fig. 1C, left middle column). Instead, with this seed, the movie condition elicited significantly more negative functional connectivity between motor and precuneus regions as compared to rest, with some more positive connectivity in the right lateral frontal cortex and scattered through some white matter regions (Fig. 1C, right middle column).

When comparing the task-evoked networks from the V1 and HV seeds (Figs. 1B and 1C, far right columns) to the corresponding resting-state networks (Figs. 1B and 1C, far left and left middle columns), we observed that these regions were more restricted using an inter-session approach than they were within-session during both the movie task and resting-state. Overall, there was little coupling of the primary and higher visual cortices to other parts of the visual system, let alone to other cortical regions.

### Whole brain patterns: task-rest FC difference versus task-evoked FC

Whole brain patterns of resting-state FC, movie FC (still containing spontaneous activity), the task-rest FC difference, and task-evoked FC were evaluated in a systematic manner using three different atlases: 1) a 17-network atlas (Yeo, et al., 2011), 2) networks obtained using spatial independent component analysis (ICA), and 3) a 246-region functional atlas (the Brainnetome Atlas) (Fan, et al., 2016). Because the ICA components used were derived in-house, we have provided them in Fig. S1; the 24 ICs that were used corresponded to 40.1% of the variance of the signal present in the concatenated data.

Similarity between the within-session resting-state and mixed (task-evoked + spontaneous) FC profiles was again made apparent, although the resting-state again showed more widely distributed FC (Fig. 2A). Within-visual functional connectivity (e.g. Vis1 to Vis2) were surprisingly weaker during the movie as compared to rest (17-network: 0 significant correlations; ICA: 3 significant correlations, t = −5.2323 to −5.2042, q = 0.0072-0.0081; 246-region: 4 significant correlations, t = −7.9544 to −6.3931, q = 0.0046-0.0196) (Figs. 2A, 3A). In contrast, task-evoked FC indicated that visual regions were positively coupled with one another due to the movie task using all three methods (17-network: 1 significant correlation, t = 7.5001, q = 0.0003; ICA: 3 significant correlations, t = 6.8589-7.9547, q = 0.0005-0.0009; 246-region: 280 significant correlations, t = 5.7474-17.2342, q = 1.5347x10^-7^-0.0291) (Fig. 2A, 3B). Further, using the 246-region atlas, every parcel within the Lateral Occipital Cortex, MedioVentral Occipital Cortex, and Fusiform Gyrus was positively connected with at least one other parcel within those regions (Fig. 3B). Overall, the within-visual correlations that dominate the task-evoked FC connectivity graph (Fig. 3B) were almost completely non-existent with respect to the task-rest FC difference (Fig. 3A).

**Figure 2.**
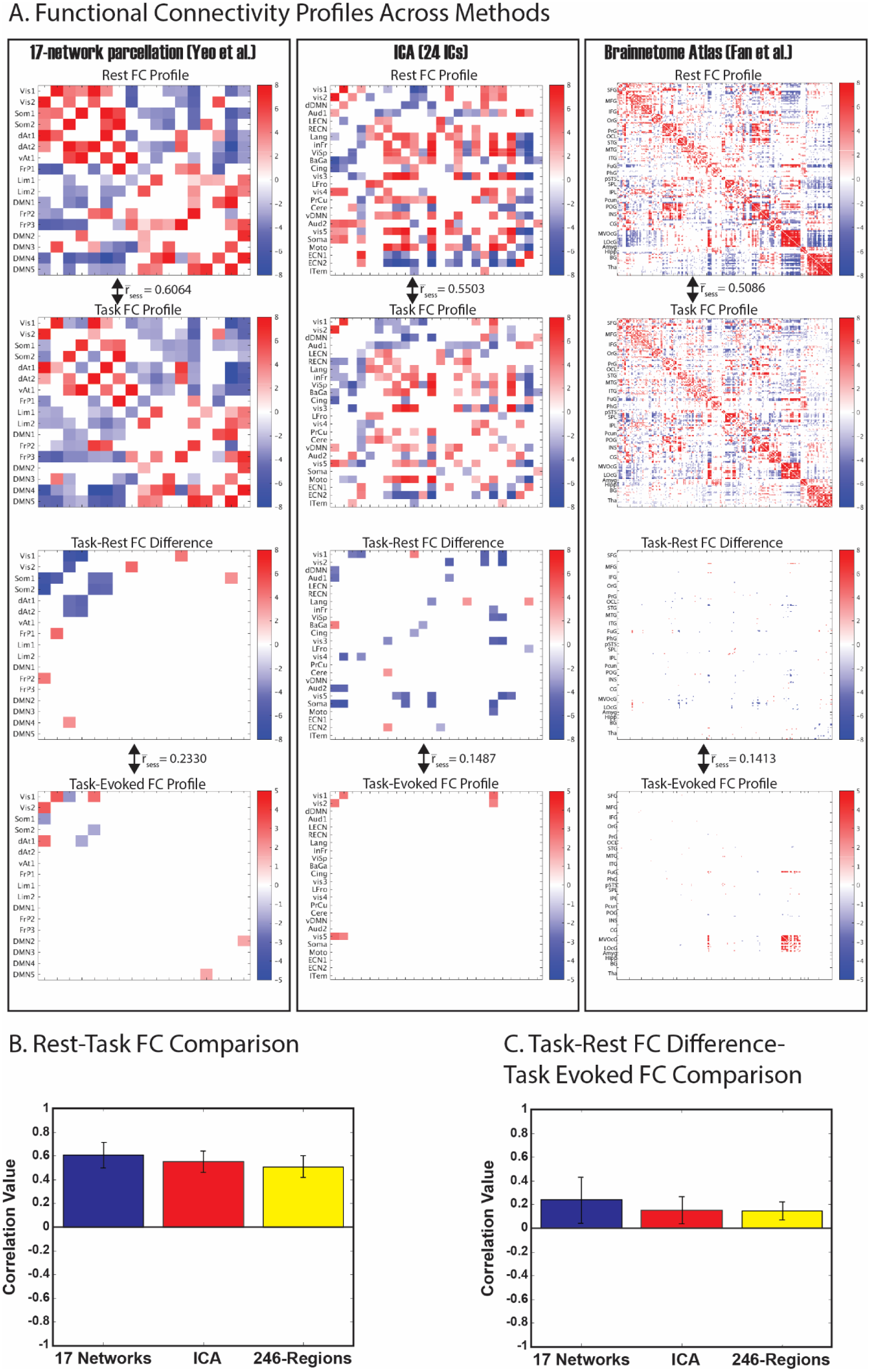
Functional Connectivity Profiles Across Methods. A) Here, we show correlation matrices corresponding to the FC profiles during resting-state (top) and the movie task (top middle), the FC difference during the movie relative to rest (bottom middle), and the task-evoked FC computed using the inter-session approach (bottom). Profiles were calculated using the Yeo et al. 17-network parcellation (left), ICA using 24 components corresponding to the canonical RSNs (middle), and the Fan et al. Brainnetome Atlas 246-region functional parcellation (right). The color bar indicates mean z-transformed cross correlation values; only significant correlations (q<0.03) are displayed. We have listed mean session-wise correlation coefficients between the resting-state and movie tasks for each of the three methods in the white space between the matrices, as well as between the task-rest FC difference and the task-evoked FC. B) The mean correlations between movie FC and rest FC are plotted on the bar graph. Error bars indicate SD. C) The mean correlations between the task-rest FC difference and task-evoked FC are plotted on the bar graph. Error bars indicate SD. Abbreviations for the Yeo parcellation: Vis1 – Visual Network 1; Vis2 – Visual Network 2; Som1 – Somatomotor Network 1; Som2 – Somatomotor Network 2; dAt1 – dorsal Attention Network 1; dAt2 – dorsal Attention Network 2; vAt1 – ventral Attention Network 1; FrP1 – Frontoparietal Network 1; Lim1 – Limbic Network 1; Lim2 – Limbic Network 2; DMN1 – Default Mode Network 1; FrP2 – Frontoparietal Network 2; FrP3 – Frontoparietal Network 3; DMN2 – Default Mode Network 2; DMN3 – Default Mode Network 3; DMN4 – Default Mode Network 4; DMN5 – Default Mode Network 5. ICA abbreviations are provided in the caption for Fig. S2, and the Brainnetome atlas abbreviations are provided in Fan, et al. (2016).

**Figure 3.**
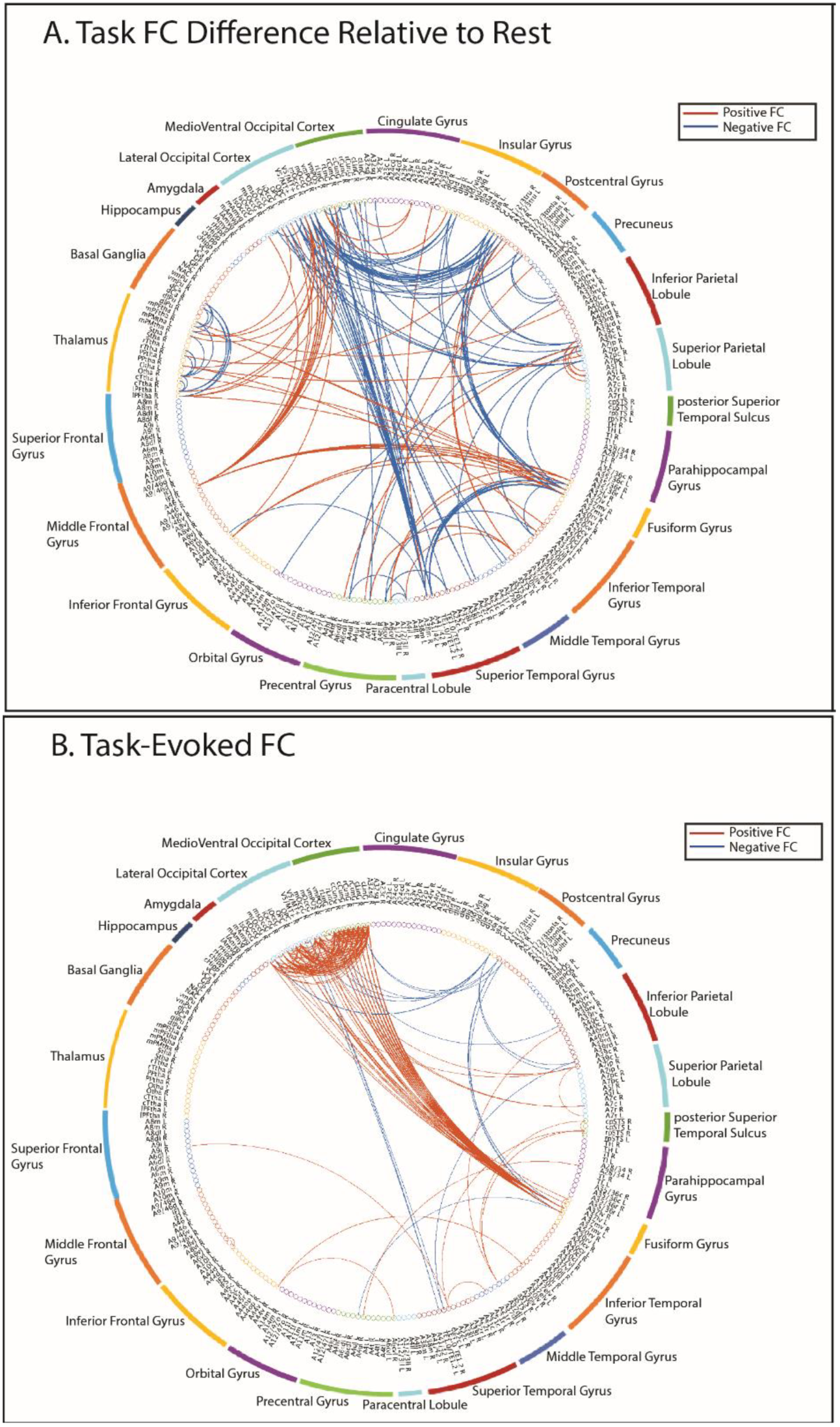
Functional Connectivity Findings: Comparing the Task-Rest FC Difference to the Task-Evoked FC. The circle graphs indicate significant FC findings (p>0.03, FDR-corrected). Abbreviations of regions are based on the Brainnetome Atlas. A) Significant Task-Rest Difference Functional Connectivity. Positive correlations during the movie relative to rest are noted with red lines; negative correlations during the movie relative to rest are noted with blue lines. B) Significant Task-Evoked Functional Connectivity. Positive correlations across two repeated viewings of the movie are denoted with red lines; negative correlations across two viewings of the movie are denoted with blue lines.

Moreover, non-visual sensory networks (e.g. somatomotor, auditory networks) exhibited weaker FC strengths with visual areas during the movie. (*Visual-Somatomotor*: 17-network: 3 significant correlations, t = −7.0622 to −4.5202, q = 0.0008-0.0250; ICA: 6 significant correlations, t = −8.0592 to −4.7280, q = 0.0003-0.0147; 246-region: 14 significant correlations, t = −7.0389 to −5.8485, q = 0.0129-0.0291; *Visual-Auditory*: 17-network: auditory cortex not assessed due to parcellation restrictions; ICA: 4 significant correlations, t = −6.5712 to −4.4074, q = 0.0012-0.0259; 246-region: 23 significantly weaker correlations, t = −11.0213 to −5.8682, q = 0.0004-0.0291.) However, when assessing task-evoked FC, we were largely unable to observe these task-rest FC differences. (*Visual-Somatomotor*: 17-network: 1 significant correlation, t = −4.9515, q = 0.0165; ICA: 0 significant correlations; 246-region: 6 significant correlations, t = −7.7223 to −5.8511, q = 0.0049-0.0246; *Visual-Auditory*: 17-network: auditory cortex not assessed, ICA: 0 significant correlations, 246-region: 3 significant correlations, t = −6.8119 to −5.8406, q = 0.0049-0.0246) (Figs. 2A, 3B).

In addition, frontoparietal networks (i.e. executive control networks) displayed stronger FC with visual regions during the movie (17-network: 4 significant correlations, t = 4.3116-4.5027, q = 0.0228-0.0281; ICA: 0 significant correlations; 246-region: 12 significant correlations, t = 5.9733-9.2107) (Figs. 2A, 3A). All significantly different correlations (positive) using the 246-region parcellation involved the right inferior frontal junction (IFJ R). Conversely, frontoparietal networks were largely not observed to have significant task-evoked FC with visual networks (17-network: 0 significant correlations; ICA: 0 significant correlations; 246-region: 4 significant correlations, t = 6.0503 – 7.2768, q = 0.0022-0.0170) (Figs. 2A, 3B). Moreover, using the 246-region atlas, the positive visual FC differences from visual areas to the inferior frontal junction (IFJ) (Fig. 3A) were not at all observed in the task-evoked FC (t = 0.0241 – 5.4687, q = 0.0472-10.8194) (Fig. 3B).

We also observed stronger visual-to-thalamus FC within-session during the movie than at rest (Fig. 3A); these differences were not observed during the task-evoked FC (Fig. 3B). (The 17-network parcellation and ICA networks did not include any thalamus-specific networks and are thus excluded from this discussion.) Eleven significant correlations were uncovered from visual regions to the thalamus when investigating the task-rest FC difference (t = 5.8489-6.4350, q = 0.0191-0.0293) (Fig. 3A). Conversely, there were zero significant task-evoked correlations between any visual and thalamus regions (t = −3.3433-4.1352, q = 0.3524-10.8733) (Fig. 3B).

Finally, by cross-correlating the resting-state and movie FC profiles, baseline connectivity patterns in the two states were highly similar, though not entirely so (mean ± SD: 17-network: r = 0.6064 ± 0.1100; ICA: r = 0.5503 ± 0.0900; 256-region: r = 0.5086 ± 0.0928) (Fig. 2B). However, the task-evoked FC and task-rest FC differences were strikingly different, and a similar correlation analysis of these two FC patterns quantitatively validated that the task-rest FC difference and task-evoked FC had very little similarity (mean ± SD: 17-network: r = 0.2330 ± 0.1923; ICA: r = 0.1487 ± 0.1120; 256-region: r = 0.1413 ± 0.0771) (Fig. 2C).

### How much of the task-rest FC difference is explained by the task-evoked activity?

After linearly regressing the task-evoked FC activity from the task-rest FC difference using 1) the Yeo et al. 17-network atlas (2011), 2) the previously obtained 24 spatial ICs, and 3) the 246-region Brainnetome Atlas (Fan, et al., 2016), we determined that the mean percent variance explained by the task-evoked activity for the 17-network atlas was 15.86 ± 3.30%, 5.19 ± 1.25% for the ICA maps, and 3.55 ± 0.73% for the 246-region atlas (all values: mean ± SEM); the mean value was calculated across sessions. Taking the mean percent variance of these three methods yielded an overall value of 8.20 ± 1.40% across both sessions and methods. Thus, only about 3-15% of the task-rest FC difference can be explained by the task-evoked activity.

## Discussion

We have shown that the difference between FC at rest and during a task, which contains an unknown mixture of task-evoked and spontaneous signals, cannot be explained by separating the task-evoked FC from the connectivity profile. The results lead to the following findings: 1) connectivity between resting-state and task states is mostly conserved; 2) during the resting-state, non-visual sensory-related functional networks (e.g. somatomotor, auditory) were more coupled to visual networks than during the movie; 3) the task-evoked FC was predominantly characterized by positive and restricted correlations among regions within the visual system, and 4) task-evoked FC accounted for only 3-15% of the FC difference between task and rest conditions. Therefore, the results suggest that the task-evoked FC and the spontaneous FC are neither linear nor additive, which was somewhat surprising to us.

### FC during a task and at rest is mostly conserved

Consistent with several prior studies (Cole, et al., 2014; Gratton, et al., 2016; Krienen, et al., 2014), we also identified a relatively high degree of similarity between the apparent FC during resting-state and the task using both seed-based and whole-brain methods (Pearson correlation values of 0.5-0.6, Fig. 2B). This is likely due to the presence of dominating spontaneous, ongoing sources in both conditions that strongly contribute to the signals correlated with one another in FC fMRI. Despite this similarity, however, we observed more widespread connectivity in the resting-state, as well as stronger within-visual coupling as compared to during the movie task.

### Apparent FC differences between rest and task are not explained by task-evoked activity

As expected, task-evoked FC was only observed within task-related, visual regions. These areas appeared to be more restricted and less coupled to other regions than in the resting-state or during the task (Fig. 1). In contrast, the connectivity differences involving visual regions between the two conditions were predominantly negative and/or not significant. Instead, we found widespread negative differences between task-related networks and non-visual sensory areas (e.g. somatomotor, auditory cortices). In addition, thalamic regions, which have not often been incorporated in analyses of FC changes, were more anti-correlated with one another and more positively correlated to portions visual cortex during the movie task. Finally, positive functional connectivity from the occipital cortex and fusiform gyrus to the inferior frontal junction (IFJ) resulted from the subtraction that also were not reproduced; functionally, the IFJ has been implicated in attentional circuits and in cognitive control (Baldauf and Desimone, 2014; Sundermann and Pfleidferer, 2012). Overall, these differences between rest and task FC were largely not represented in the task-evoked FC patterns.

The fact that the task-evoked FC did not reveal the difference between the FC during the task and the FC at rest (i.e. spontaneous FC) suggests that correlations in ongoing, spontaneous activity are driving this difference. Therefore, it is likely that this intrinsic activity drives the coupling of task-evoked networks to other regions.

### Rest and task correlations negatively interact

The task-evoked FC explained less than 15% of the FC differences between the task and resting-state. Therefore, it seems that the task-evoked FC and spontaneous FC are neither independent nor linearly additive. Beyond this, however, we would like to tease apart the nature of the rest-task interaction: is the task suppressing spontaneous activity or amplifying it? Our observations that the movie-watching task reduced the extent and strength of FC suggest that the task suppresses spontaneous activity.

He (2013) and several others (Bianciardi, et al., 2009; Churchland, et al., 2010; Monier, et al., 2003; Ponce-Alvarez, et al., 2013), also suggest a negative task-rest interaction. Initial evidence suggests that this negative interaction may help facilitate the task execution (Boly, et al., 2007; Deneux and Grinvald, 2017; Hesselmann, et al., 2008) (see Northoff, et al. (2010) and Ferezou and Deneux (2017) for review), or may increase with task difficulty (Garrett, et al., 2014; Szostakiwskyj, et al., 2017). As such, it may bear functional significance.

The negative task-rest interaction may or may not hold true for all tasks. Passive versus active task engagement may not equally affect spontaneous signals (Broday-Divir, et al., 2017; Ferezou, et al., 2006; Otazu, et al., 2009). Crochet and Petersen (2006) found that active and conscious engagement in a task gave rise to more desynchronization of ongoing activity than passive or conscious states (e.g. in the anesthetized states). In our natural vision task, subjects actively engaged in the movie with free eye movement. Speculatively, cognitively engaging in the task itself, rather than simply having a visual experience, explains the nonlinear interaction between spontaneous and evoked functional connectivity. However, this remains to be tested.

Using natural vision, we noticed that the suppression of spontaneous correlations during the task was not consistent throughout the brain. The greatest magnitude of this change was within the components of the visual system; these regions exhibited the greatest dissimilarity between task-evoked FC and the apparent FC difference between the movie and resting-state conditions. These findings may be mediated simply by 1) reduced spontaneous activations in visual areas relative to other regions, or 2) by a reduced synchrony of cortical oscillations in task-related regions. In EEG, alpha band oscillations are postulated to stem from the rhythmic fluctuations of inhibitory neurons, and engaging in certain tasks such as eye-opening, desynchronizes the alpha-band power (see Klimesch, et al. (2007) for review). Other reports relate resting-state inhibitory neurotransmitter concentrations, such as GABA (Muthukumaraswamy, et al., 2009; Northoff, et al., 2007) or anesthetics thought to modulate GABA (Maandag, et al., 2007), to task-induced changes in specific regions. Here, we cannot disentangle whether location differences in spontaneous FC suppression are mediated by region-specific reduced activations or de-coupling of neuronal oscillations, but this is certainly an area for future investigation.

### Methodological Considerations

Indeed, naturalistic stimuli (Hasson, et al., 2004) are of particular significance in studies of rest-task interaction. Natural stimuli provide a rich behavioral context reflective of the activities of daily life (e.g. viewing natural scenes with sharp, moving edges or engaging in conversation) that unfold over relatively long time scales (Hasson, et al., 2010). It has experimentally been proven that neural responses to naturalistic stimuli are reliable and widespread (Hasson, et al., 2010; Jääskeläinen, et al., 2008; McMahon, et al., 2015; Mukamel, et al., 2005), and the connectivity patterns that appear during naturalistic activations better reflect spontaneously emerging patterns in the resting-state as compared to controlled, artificially designed stimuli (Wilf, et al., 2017). Our group has shown that by spatiotemporally scrambling the natural stimulus, widely distributed and highly reproducible fMRI responses could not be reproduced without the high-level natural content of the movie; low-level visual features alone significantly reduced the degree and extent of reproducible responses (Lu, et al., 2016). Therefore, naturalistic visual stimuli provide rich task-evoked information about neural dynamics as compared to more traditional psychophysical stimuli (e.g. Gabor filters).

Optimally isolating the task-evoked activity is important for studies of rest-task interaction. One way of reducing the variability present in fMRI signals is through temporal averaging; however, a very large number of subjects and/or sessions is needed to achieve appropriate statistical power. Even with a great number samples, the efficacy of simple averaging in removing spontaneous activity has limitations (Henriksson, et al., 2015; Kim, et al., 2017). An earlier approach uses the general linear model (GLM) to construct a trial-to-trial series of activation parameters (β) for each voxel that can be cross-correlated (Mennes, et al., 2013; Rissman, et al., 2004); however, whether this method more effectively removes intrinsic activity than inter-session and inter-subject approaches has yet to be shown. Finally, between inter-session (i.e. “intra-subject”) and inter-subject approaches, inter-session correlations have shown enhanced reproducibility. (Henriksson, et al., 2015; Lu, et al., 2016).

Inter-session and inter-subject correlation methods have been understudied in neuroimaging, and new studies using these methods provide an additional vantage point from which we may learn about the brain. In this work, our focus was on whether the difference between the resting-state and the mixed FC observed during the task reflected the task-evoked FC. It did not, but we shed light on a suppression of correlations of spontaneous activity that occurs to facilitate a task. However, a consensus regarding this phenomenon still needs to be formed for additional researchers to fully disentangle its origins and purpose.

## Acknowledgments

The research was supported in part by NIH R01MH104402 (Z. Liu, PI) and a predoctoral fellowship awarded to Lauren Lynch as supported by TL1 TR001107 and UL1 TR001108 (A. Shekhar, PI) from the National Institutes of Health, National Center for Advancing Translational Sciences, Clinical and Translational Sciences Award. AJS acknowledges support from NIH grants P30 AG010133 and R01 AG019771.

---

**Figure S1.**
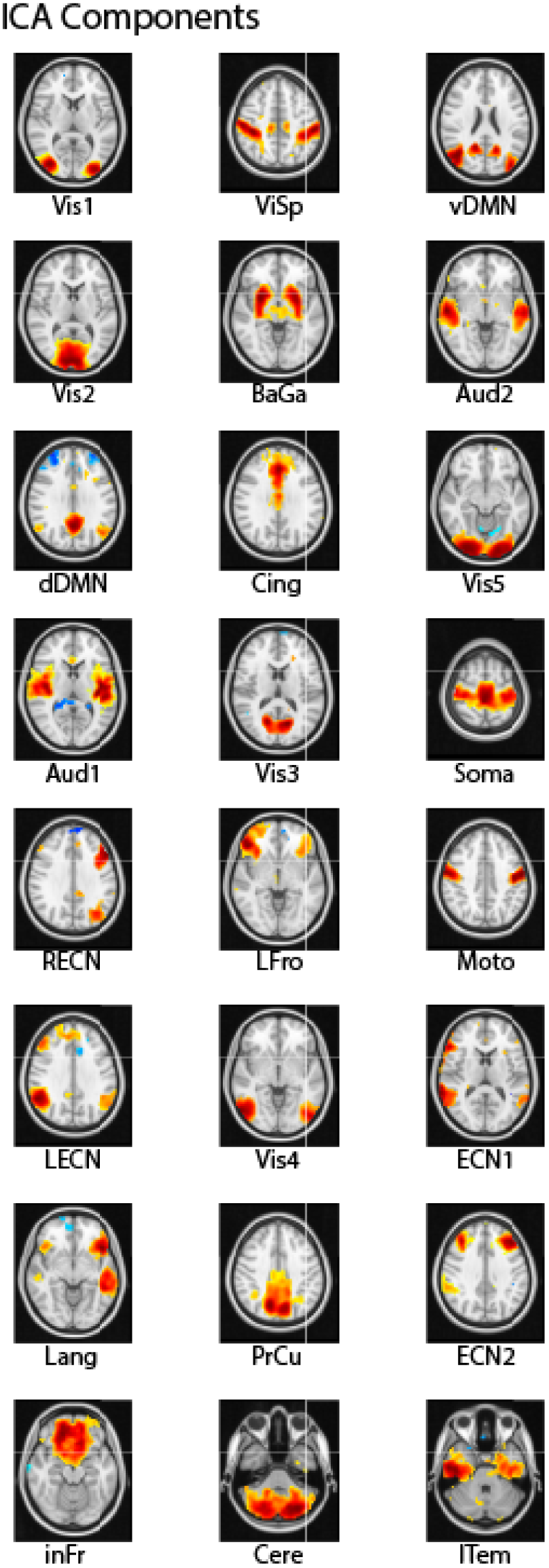
ICA Components. Maps obtained using group-level spatial ICA. The thresholding for display purposes only was determined according to the voxel-wise posterior probability equal to 0.6, per a Gaussian Mixture Model; ICA maps used in any calculations were not thresholded. Abbreviations from top-to-bottom, left-to-right are as follows: Visual Network 1 (Vis1), Visual Network 2 (Vis2), dorsal Default Mode Network (dDMN), Auditory Network 1, Right Executive Control Network (RECN), Left Executive Control Network (LECN), Language Network (Lang), inferior Frontal Network (inFr), Visual-Spatial Network (ViSp), Basal Ganglia (BaGa), Cingulate Network (Cing), Visual Network 3 (Vis3), Lateral Frontal Network (LFro),Visual Network 4 (Vis4), Precuneus (PrCu), Cerebellum Network 1 (Cer1), ventral Default Mode Network (vDMN), Auditory Network 2 (Aud2), Visual Network 5 (Vis5), Somatosensory Network (Soma), Motor Network (Moto), Executive Control Network 1 (ECN1), Executive Control Network 2 (ECN2), and Cerebellum Network 2 (Cer2).

**Figure S2.**
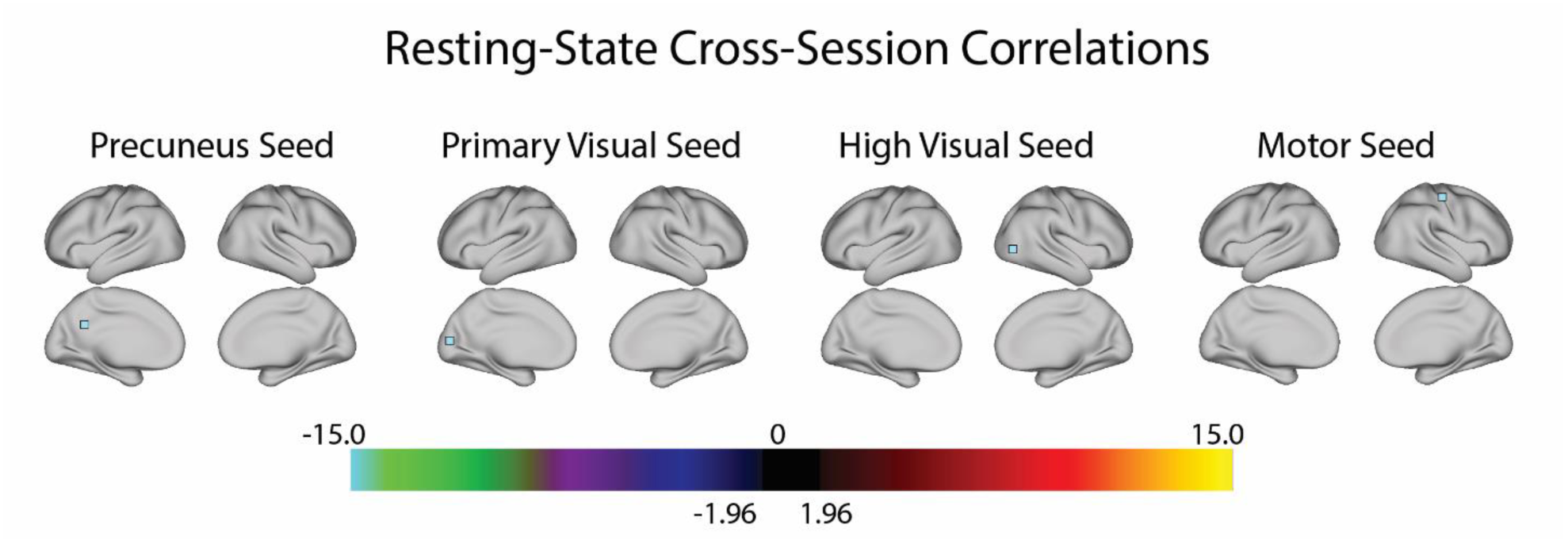
Resting-state Inter-session Correlations. By showing that there are no significant voxels correlated to the seed voxel across two sessions of the same stimulus, we demonstrate the efficacy of inter-session correlations in isolating task-evoked activity. The seed voxels were the same as in Fig. 1 and were derived from the precuneal (left), B) primary visual (left middle), C) high visual (right middle), and D) motor cortices (right), respectively. The color bar indicates z-transformed cross correlation values.

## Footnotes

1 Downloaded 18 May 2017

2 Downloaded 21 July 2017

3 Downloaded 13 September 2017

## References

Arfanakis, K., Cordes, D., Haughton, V.M., Moritz, C.H., Quigley, M.A., Meyerand, M.E. (2000) Combining independent component analysis and correlation analysis to probe interregional connectivity in fMRI task activation datasets. Magnetic Resonance Imaging, 18:921–930.

Arieli, A., Sterkin, A., Grinvald, A., Aertsen, A. (1996) Dynamics of ongoing activity: explanation of the large variability in evoked cortical responses. Science, 273:1868–1871.

Azouz, R., Gray, C.M. (1999) Cellular mechanisms contributing to response variability of cortical neurons in vivo. The Journal of Neuroscience, 19:2209–2223.

Baldauf, D., Desimone, R. (2014) Neural Mechanisms of Object-Based Attention. Science, 344:424–427.

Becker, R., Reinacher, M., Freyer, F., Villringer, A., Ritter, P. (2011) How Ongoing Neuronal Oscillations Account for Evoked fMRI Variability. Journal of Neuroscience, 31:11016–11027.

Beckmann, C.F., DeLuca, M., Devlin, J.T., Smith, S.M. (2005) Investigations into resting-state connectivity using independent component analysis. Philosophical transactions of the Royal Society of London. Series B, Biological sciences, 360:1001–1013.

Bell, a.J., Sejnowski, T.J. (1995) An information-maximization approach to blind separation and blind deconvolution. Neural computation, 7:1129–1159.

Bianciardi, M., Fukunaga, M., Gelderen, P.V., Horovitz, S.G., De, J.A., Duyn, J.H. (2009) Modulation of spontaneous fMRI activity in human visual cortex by behavioral state. NeuroImage, 45:160–168.

Boly, M., Balteau, E., Schnakers, C., Degueldre, C., Moonen, G., Luxen, A., Phillips, C., Peigneux, P. (2007) Baseline brain activity fluctuations predict somatosensory perception in humans. Proceedings of the National Academy of Sciences of the United States of America, 104:12187–12192.

Borg-Graham, L.J., Monier, C., Fregnac, Y. (1998) Visual input evokes transient and strong shunting inhibition in visual cortical neurons. Nature, 393:369–373.

Brainard, D.H. (1997) The Psychophysics Toolbox. Spatial Vision, 10:433–436.

Broday-Divir, R., Grossman, S., Furman-Haran, E., Malach, R. (2017) Quenching of spontaneous fluctuations by attention in human visual cortex. Neuroimage.

Buckner, R.L., Krienen, F.M., Yeo, T.B.T. (2013) Opportunities and limitations of intrinsic functional connectivity MRI. Nature Reviews Neuroscience, 16:832–837.

Chang, C., Glover, G. (2010) Time-frequency dynamics of resting-state brain connectivity measured with fMRI. NeuroImage, 50:81–98.

Churchland, M.M., Yu, B.M., Cunningham, J.P., Sugrue, L.P., Cohen, M.R., Corrado, G.S., Newsome, W.T., Clark, A.M., Hosseini, P., Scott, B.B., Bradley, D.C., Smith, M.A., Kohn, A., Movshon, J.A., Armstrong, K.M., Moore, T., Chang, S.W., Snyder, L.H., Lisberger, S.G., Priebe, N.J., Finn, I.M., Ferster, D., Ryu, S.I., Santhanam, G., Sahani, M., Shenoy, K.V. (2010) Stimulus onset quenches neural variability: a widespread cortical phenomenon. Nature Neuroscience, 13:369–378.

Cole, M.W., Bassett, D.S., Power, J.D., Braver, T.S., Petersen, S.E. (2014) Intrinsic and task-evoked network architectures of the human brain. Neuron, 83:238–251.

Cox, R.W. (1996) AFNI: Software for Analysis and Visualization of Functional Magnetic Resonance Neuroimages. Computers and Biomedical Research, 29:162–173.

Crochet, S., Petersen, C.C. (2006) Correlating whisker behavior with membrane potential in barrel cortex of awake mice. Nat Neurosci, 9:608–10.

De Luca, M., Beckmann, C.F., De Stefano, N., Matthews, P.M., Smith, S.M. (2006) fMRI resting state networks define distinct modes of long-distance interactions in the human brain. NeuroImage, 29:1359–67.

Deneux, T., Grinvald, A. (2017) Milliseconds of Sensory Input Abruptly Modulate the Dynamics of Cortical States for Seconds. Cereb Cortex, 27:4549–4563.

Fair, D.A., Schlaggar, B.L., Cohen, A.L., Miezin, F.M., Dosenbach, N.U.F., Wenger, K.K., Fox, M.D., Snyder, A.Z., Raichle, M.E., Petersen, S.E. (2007) A method for using blocked and event-related fMRI data to study “resting state” functional connectivity. NeuroImage, 35:396–405.

Fan, L., Li, H., Zhuo, J., Zhang, Y., Wang, J., Chen, L., Yang, Z., Chu, C., Xie, S., Laird, A.R., Fox, P.T., Eickhoff, S.B., Yu, C., Jiang, T. (2016) The Human Brainnetome Atlas: A New Brain Atlas Based on Connectional Architecture. Cereb Cortex, 26:3508–26.

Ferezou, I., Bolea, S., Petersen, C.C. (2006) Visualizing the cortical representation of whisker touch: voltage-sensitive dye imaging in freely moving mice. Neuron, 50:617–29.

Ferezou, I., Deneux, T. (2017) Review: How do spontaneous and sensory-evoked activities interact? Neurophotonics, 4:031221.

Finn, I.M., Priebe, N.J., Ferster, D. (2007) The emergence of contrast-invariant orientation tuning in simple cells of cat visual cortex. Neuron, 54:137–152.

Fox, M.D., Snyder, A.Z., Zacks, J.M., Raichle, M.E. (2006) Coherent spontaneous activity accounts for trial-to-trial variability in human evoked brain responses. Nature Neuroscience, 9:23–5.

Garrett, D.D., McIntosh, A.R., Grady, C.L. (2014) Brain signal variability is parametrically modifiable. Cerebral Cortex, 24:2931–2940.

Gonzalez-Castillo, J., Hoy, C.W., Handwerker, D.a., Robinson, M.E., Buchanan, L.C., Saad, Z.S., Bandettini, P.A. (2015) Tracking ongoing cognition in individuals using brief, whole-brain functional connectivity patterns. Proceedings of the National Academy of Sciences, 112:8762–8767.

Gratton, C., Laumann, T.O., Gordan, E.M., Adeyemo, B., Petersen, S.E. (2016) Evidence for two independent factors that modify brain networks to meet task goals. Cell Report, 16:338–348.

Greve, D.N., Fischl, B. (2009) Accurate and robust brain image alignment using boundary-based registration. Neuroimage, 48:63–72.

Harrison, B.J., Pujol, J., López-Solà, M., Hernández-Ribas, R., Deus, J., Ortiz, H., Soriano-Mas, C., Yücel, M., Pantelis, C., Cardoner, N. (2008) Consistency and functional specialization in the default mode brain network. Proceedings of the National Academy of Sciences of the United States of America, 105:9781–9786.

Hasson, U., Malach, R., Heeger, D.J. (2010) Reliability of cortical activity during natural stimulation. Trends in Cognitive Sciences, 14:40–48.

Hasson, U., Nir, Y., Levy, I., Fuhrmann, G., Malach, R. (2004) Intersubject synchronization of cortical activity during natural vision. Science, 303:1634–1640.

He, B.J. (2013) Spontaneous and task-evoked brain activity negatively interact. Journal of Neuroscience, 33:4672–4682.

Henriksson, L., Khaligh-Razavi, S.M., Kay, K., Kriegeskorte, N. (2015) Visual representations are dominated by intrinsic fluctuations correlated between areas. NeuroImage, 114:275–286.

Hesselmann, G., Kell, C.A., Eger, E., Kleinschmidt, A. (2008) Spontaneous local variations in ongoing neural activity bias perceptual decisions. Proceedings of the National Academy of Sciences of the United States of America, 105:10984–10989.

Horovitz, S.G., Fukunaga, M., De Zwart, J.A., Van Gelderen, P., Fulton, S.C., Balkin, T.J., Duyn, J.H. (2008) Low frequency BOLD fluctuations during resting wakefulness and light sleep: A simultaneous EEG-fMRI study. Human Brain Mapping, 29:671–682.

Hutchison, R.M., Womelsdorf, T., Allen, E.A., Bandettini, P.A., Calhoun, V.D., Corbetta, M., Penna, S.D., Duyn, J.H., Glover, G.H., Gonzalez-Castillo, J., Handwerker, D.A., Keilholz, S., Kiviniemi, V., Leopold, D.A., de Pasquale, F., Sporns, O., Walter, M., Chang, C. (2013) Dynamic functional connectivity: Promise, issues, and interpretations. NeuroImage, 80:1–43.

Jääskeläinen, I.P., Koskentalo, K., Balk, M.H., Autti, T., Kauramäki, J., Pomren, C., Sams, M. (2008) Inter-subject synchronization of prefrontal cortex hemodynamic activity during natural viewing. The Open Neuroimaging Journal, 2:14–19.

Kim, D., Kay, K., Shulman, G.L., Corbetta, M. (2017) A New Modular Brain Organization of the BOLD Signal during Natural Vision. Cerebral Cortex:1–17.

Klimesch, W., Sauseng, P., Hanslmayr, S. (2007) EEG alpha oscillations: the inhibition-timing hypothesis. Brain Res Rev, 53:63–88.

Krienen, F.M., Yeo, B.T.T., Buckner, R.L., Buckner, R.L. (2014) Reconfigurable task-dependent functional coupling modes cluster around a core functional architecture. Philosophical transactions of the Royal Society B.

Lu, K.H., Hung, S.C., Wen, H., Marussich, L., Liu, Z. (2016) Influences of high-level features, gaze, and scene transitions on the reliability of BOLD responses to natural movie stimuli. PLoS ONE, 11:1–19.

Maandag, N.J.G., Coman, D., Sanganahalli, B.G., Herman, P., Smith, A.J., Blumenfeld, H., Shulman, R.G., Hyder, F. (2007) Energetics of neuronal signaling and fMRI activity. Proceedings of the National Academy of Sciences of the United States of America, 104:20546–20551.

Mäkinen, V., Tiitinen, H., May, P. (2005) Auditory event-related responses are generated independently of ongoing brain activity. NeuroImage, 24:961–968.

Marussich, L., Lu, K.-H., Wen, H., Liu, Z. (2017) Mapping White-Matter Functional Organization at Rest and during Naturalistic Visual Perception. NeuroImage, 146:1128–1141.

McMahon, D.B.T., Russ, B.E., Elnaiem, H.D., Kurnikova, a.I., Leopold, D.a. (2015) Single-Unit Activity during Natural Vision: Diversity, Consistency, and Spatial Sensitivity among AF Face Patch Neurons. Journal of Neuroscience, 35:5537–5548.

Mennes, M., Kelly, C., Colcombe, S., Xavier Castellanos, F., Milham, M.P. (2013) The extrinsic and intrinsic functional architectures of the human brain are not equivalent. Cerebral Cortex, 23:223–229.

Monier, C., Chavane, F., Baudot, P., Graham, L.J., Frégnac, Y. (2003) Orientation and direction selectivity of synaptic inputs in visual cortical neurons: A diversity of combinations produces spike tuning. Neuron, 37:663–680.

Mukamel, R., Gelbard, H., Arieli, A., Hasson, U., Fried, I., Malach, R. (2005) Coupling Between Neuronal Firing, Field Potentials, and fMRI in Human Auditory Cortex. Science, 309:951–954.

Muthukumaraswamy, S.D., Edden, R.A.E., Jones, D.K., Swettenham, J.B., Singh, K.D. (2009) Resting GABA concentration predicts peak gamma frequency and fMRI amplitude in response to visual stimulation in humans. Proceedings of the National Academy of Sciences of the United States of America, 106:8356–8361.

Northoff, G., Qin, P., Nakao, T. (2010) Rest-stimulus interaction in the brain: a review. Trends Neurosci, 33:277–84.

Northoff, G., Walter, M., Schulte, R.F., Beck, J., Dydak, U., Henning, A., Boeker, H., Grimm, S., Boesiger, P. (2007) GABA concentrations in the human anterior cingulate cortex predict negative BOLD responses in fMRI. Nat Neurosci, 10:1515–7.

Oram, M.W. (2011) Visual Stimulation Decorrelates Neuronal Activity. Journal of Neurophysiology, 105:942–957.

Otazu, G.H., Tai, L.H., Yang, Y., Zador, A.M. (2009) Engaging in an auditory task suppresses responses in auditory cortex. Nat Neurosci, 12:646–54.

Pelli, D.G. (1997) The VideoToolbox software for visual psychophysics: Transforming numbers into movies. Spatial Vision, 10:437–442.

Ponce-Alvarez, A., Thiele, A., Albright, T.D., Stoner, G.R., Deco, G. (2013) Stimulus-dependent variability and noise correlations in cortical MT neurons. Proceedings of the National Academy of Sciences, 110:13162–13167.

Power, J.D., Cohen, A.L., Nelson, S.M., Wig, G.S., Barnes, K.A., Church, J.A., Vogel, A.C., Laumann, T.O., Miezin, F.M., Schlaggar, B.L., Petersen, S.E. (2011) Functional Network Organization of the Human Brain. Neuron, 72:665–678.

Rehme, A.K., Eickhoff, S.B., Grefkes, C. (2013) State-dependent differences between functional and effective connectivity of the human cortical motor system. NeuroImage, 67:237–246.

Rissman, J., Gazzaley, A., D’Esposito, M. (2004) Measuring functional connectivity during distinct stages of a cognitive task. NeuroImage, 23:752–763.

Saka, M., Berwick, J., Jones, M. (2010) Linear superposition of sensory-evoked and ongoing cortical hemodynamics. Frontiers in neuroenergetics, 2:1–13.

Sepulcre, J., Liu, H., Talukdar, T., Martincorena, I.i., Thomas Yeo, B.T., Buckner, R.L. (2010) The organization of local and distant functional connectivity in the human brain. PLoS Computational Biology, 6:1–15.

Shirer, W.R., Ryali, S., Rykhlevskaia, E., Menon, V., Greicius, M.D. (2012) Decoding Subject-Driven Cognitive States with Whole-Brain Connectivity Patterns. Cerebral Cortex, 22:158–165.

Simony, E., Honey, C.J., Chen, J., Lositsky, O., Yeshurun, Y., Wiesel, A., Hasson, U. (2016) Dynamical reconfiguration of the default mode network during narrative comprehension. Nature Communications, 7:1–13.

Smith, S.M., Jenkinson, M., Woolrich, M.W., Beckmann, C.F., Behrens, T.E.J., Johansen-Berg, H., Bannister, P.R., De Luca, M., Drobnjak, I., Flitney, D.E., Niazy, R.K., Saunders, J., Vickers, J., Zhang, Y., De Stefano, N., Brady, J.M., Matthews, P.M. (2004) Advances in functional and structural MR image analysis and implementation as FSL. NeuroImage, 23:208–219.

Sundermann, B., Pfleidferer, B. (2012) Functional connectivity profile of the human inferior frontal junction: involvement in a cognitive control network. BMC Neuroscience, 3.

Szostakiwskyj, J.M.H., Willatt, S.E., Cortese, F., Protzner, A.B. (2017) The modulation of EEG variability between internally- and externally-driven cognitive states varies with maturation and task performance. Plos One, 12:e0181894.

Tsodyks, M., Kenet, T., Grinvald, A., Arieli, A. (1999) Linking spontaneous activity of single cortical neurons and the underlying functional architecture. Science (New York, N.Y.), 286:1943–1946.

Van Dijk, K.R.a., Hedden, T., Venkataraman, A., Evans, K.C., Lazar, S.W., Buckner, R.L. (2010) Intrinsic Functional Connectivity As a Tool For Human Connectomics: Theory, Properties, and Optimization. Journal of Neurophysiology, 103:297–321.

Vincent, J.L., Patel, G.H., Fox, M.D., Snyder, a.Z., Baker, J.T., Van Essen, D.C., Zempel, J.M., Snyder, L.H., Corbetta, M., Raichle, M.E. (2007) Intrinsic functional architecture in the anaesthetized monkey brain. Nature, 447:83–86.

Wilf, M., Strappini, F., Golan, T., Hahamy, A., Harel, M., Malach, R. (2017) Spontaneously emerging patterns in human visual cortex reflect responses to naturalistic sensory stimuli. Cerebral Cortex, 27:750–763.

Wong, C.W., Olafsson, V., Tal, O., Liu, T.T. (2013) The amplitude of the resting-state fMRI global signal is related to EEG vigilance measures. Neuroimage, 83:983–90.

Yeo, B.T.T., Krienen, F.M., Sepulcre, J., Sabuncu, M.R., Lashkari, D., Hollinshead, M., Roffman, J.L., Smoller, J.W., Zollei, L., Polimeni, J.R., Fischl, B., Liu, H., Buckner, R.L. (2011) The organization of the human cerebral cortex estimated by intrinsic functional connectivity. Journal of Neurophysiology, 106:1125–1165.

